# Reduced ornamentation became elaborated in benign environments in a bird species

**DOI:** 10.1101/2023.08.10.552863

**Authors:** Masaru Hasegawa, Emi Arai, Takahiro Kato

## Abstract

Many empirical studies have focused on highly-ornamented species to identify ecological factors that maintain the ornamentation or favour its exaggeration. By contrast, although reduction or loss of ornamentation appears to be widespread, species with reduced/lost ornamentation and its relationship with ecological factors are rarely focused on. Here, based on data collected over four years, we studied outermost tail feather length, i.e. a well-known sexual ornamentation in this clade, in relation to roosting location in the Pacific swallow *Hirundo tahitica* during winter. In contrast to congeners, this species has inconspicuous tail ornamentation, i.e. very shallowly forked tails with vestigial streamers, providing a rare opportunity to study the ecological factors driving reduced ornamentation. We found that Pacific swallows mainly roost in old nests under bridges over rivers, which resemble their original roosting sites, but some roost in old nests under the eaves of houses above the ground. Individuals roosting under the eaves of houses had significantly longer outermost (but not central) tail feathers than those roosting under bridges. Individuals roosting under the eaves of houses were heavier and showed lower physiological stress, and thus might better endure the maintenance cost, favouring ornament elaboration. Because we controlled for the effects of sex and age, these factors would not confound the observed pattern. Reduced ornamentation, as found in Pacific swallows roosting under their original roosting sites, could become elaborated in benign environments (i.e., under eaves, here), stressing the importance of balance between the costs and benefits of ornamentation.

## INTRODUCTION

Animals sometimes exhibit conspicuous ornaments, for example, elongated tails and colourful plumage in birds, that appear to have a rather negative effect on survivorship (reviewed in Andersson 1994). Many empirical studies have focused on highly-ornamented species and intraspecific patterns to identify evolutionary and ecological factors that maintain ornamentation or favour its exaggeration. By contrast, although reduction or loss of ornamentation appears to be widespread, poorly-ornamented species receive little attention (reviewed in Wiens 2001; Heinen-Kay and Zuk 2019). Given that sexual selection is a strong force preventing ornament loss at least in theory (e.g. Weigel et al. 2015), empirical studies should focus on factors that might balance with sexual selection in such species. Ornaments should be maintained because of the benefits and costs of their expression (Heinen-Kay and Zuk 2019), except when they are selectively neutral (i.e. an obsolete signal, which means no longer a signal in the macroevolutionary timescale; van Doorn and Weissing 2004).

Anthropogenic environments, which have expanded rapidly worldwide (Marzluff 2001), provide a suitable opportunity to study the function of presumably sexually selected traits, including reduced ornaments. Such environments contain new ecological niches, in which the benefits and costs of a given ornament differ from those in the ancestral habitat (e.g. see Thompson et al. 2019; Otto 2018; Sepp et al. 2020; Cronin et al. 2022 for recent reviews). For example, the túngara frog *Engystomops pustulosus* increased its song conspicuousness in urban habitats due to the reduced predation pressure (Halfwerk et al. 2018). Likewise, such environments can provide animals with unforeseen favourable conditions under which the benefits of ornaments outweigh their costs, which would allow reduced ornaments to become exaggerated again.

The Pacific swallow *Hirundo tahitica* is a congener of a well-known model species of sexual selection, the Barn swallow *H. rustica* (Turner and Rose 1994; Turner 2006), in which sexual selection for long tails has been repeatedly demonstrated (reviewed in Romano et al. 2017). They are socially monogamous non-migratory birds and display biparental provisioning but show female-only incubation, in which opportunity for extra-pair matings should be high (see above; Turner & Rose 1994). Unlike barn swallows and many other congeners, which have deeply forked tails with long tail streamers, the Pacific swallow has evolved a very shallowly forked tail with vestigial tail streamers. In other words, they have lost the long tail, as revealed by the estimated evolutionary trajectories of tails (Fig. 1; Turner and Rose 1994). Although a long outermost tail feather, or “streamer”, was once considered an aerodynamic device to enhance manoeuvrability, which might increase foraging efficiency (e.g., Buchanan & Evans 2000), a series of recent macroevolutionary studies suggest that long outermost tail feathers and hence forked tails evolve and are maintained not by efficient aerial foraging but by sexual selection (e.g., Hasegawa 2022, 2023; Hasegawa and Arai 2018, 2020a,b, 2022): hirundine species with limited extrapair opportunities are more likely to lose their long, sexually dimorphic tails. The series of studies also suggest that not a change in the direction of sexual selection such as selection for and against long tails, but the intensity of sexual selection for long tails explains the variety of tail lengths in this clade.

**Figure 1.**
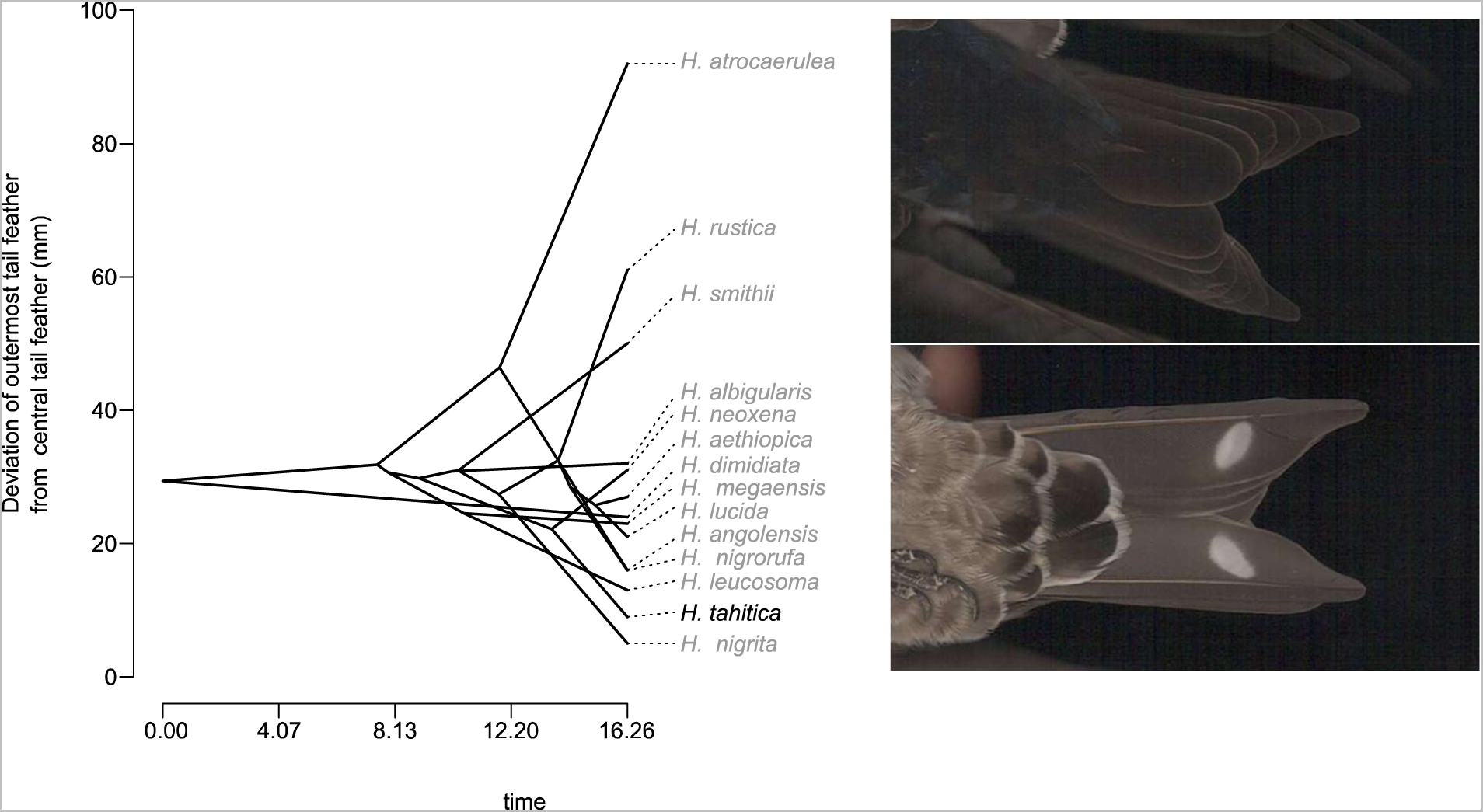
Phenogram of fork tail depth, i.e. deviation of the outermost tail feather from the central tail feather, in the genus *Hirundo*. The phylogeny and dataset were obtained from birdtree.org and Turner and Rose (1994), respectively (note that *H. neoxena*, *H. aethiopiaca*, *H. megaensis*, *H. lucida*, *H. angolensis*, *H. leucosoma*, *H. tahitica*, and *H. nigrita* are resident species). The phenogram was computed using the function “phenogram” in the R package “phytools” (Revell 2012). Note that the estimated evolutionary trajectory to *H. tahitica* shows a negative trend, indicating that their outermost tail feathers have shorten throughout time. An example of tail feathers of the Pacific swallow *H. tahitica* is shown next to the phenogram. Upper panel, dorsal side of the tail; lower panel, ventral side of the tail. Both photographs are of the same individual. Note that the outermost tail feather has a short “streamer” part and is only slightly longer than the central tail feathers, producing a shallowly forked tail.

In the focal species, Pacific swallows, males have statistically longer outermost tail feathers than females, but the sex difference is negligible (ca. 2% longer in males: Hasegawa and Arai 2017a, Hasegawa et al. 2019); this differs from the barn swallow, which exhibits marked sexual dimorphism in tail length (ca. 20% longer in males: Turner and Rose 1994; also see Hasegawa et al. 2010 for Japanese subspecies). These findings, together with macroevolutionary evidence, suggest reduced sexual selection for a long outermost tail feather, rather than a lack of sexual selection or sexual selection for a short outermost tail feather in the Pacific swallow. This sharply contrasts with other bird species that showed “reversed” sexual dimorphism in tail length (e.g., Balmford et al. 2000; Swaddle et al. 2000), although behavioural tests of mate choice have yet to be conducted in this species. A natural experiment has recently shown that long tails have a selective disadvantage during foraging, at least during severe winter weather (Hasegawa and Arai 2017a, Hasegawa et al. 2019), suggesting that even short tails of Pacific swallows are not neutral but are maintained as the consequence of balanced selection. The Pacific swallow is a non-migratory species and thus cannot avoid inclement weather through long-distance dispersal; therefore, its ornamentation would strongly be affected by the local environment (Hasegawa et al. 2016, 2019). Nonetheless, whether their tails would be elaborated in benign environments, which is predicted if ornament expression reflects the costs and benefits of ornamentation (i.e., tails still functioning as a signal even in this species with reduced ornamentation), rather than being a selectively neutral, obsolete signal (i.e., not used as a signal in this species), remains unclear.

Here, we examined outermost tail feather length in the Pacific swallow in relation to the local environment on Amami Oshima Island. For this purpose, we focused on the locations of winter roosts. Although previous sexual selection studies have often overlooked the wintering period, it should be noted that, after moulting in the late summer, only birds that survive the wintering period can reproduce and thus the survival cost and reproductive advantage are not additive, but multiplicative (Fromhage and Henshaw 2022). During winter, Pacific swallows typically roost in old nests under bridges over rivers, perhaps to avoid predation by terrestrial animals, but some roost in old nests under the eaves of buildings, such as houses and apartment buildings, located over the ground (Fig. 2). An important difference between the two types of roost is that the former, which resemble their natural breeding/roosting sites, i.e. ledges of cliffs, which are “usually over or close to water” in this species (Turner and Rose 1994), are much cooler than the latter due to wind, water vapour, and a lack of heat insulation (e.g. Murakami et al. 1991; Manteghi et al. 2015; also see Diamond and Martin 2020; Cronin et al. 2022 for the urban heat island effect on human residential areas). The difference between the two types of roosting location is particularly notable at night, as solar heat that accumulates in buildings during the day is released at night, reducing diel temperature variation in human residential areas (Diamond and Martin 2020; Cronin et al. 2022). Therefore, Pacific swallows roosting under bridges require greater investment in somatic maintenance compared to those roosting in human residential areas away from water. Longer tails require a higher maintenance cost because they impair foraging ability (e.g. Hasegawa and Arai 2017a, 2020a,b, 2022; Hasegawa et al. 2019), which might explain the short tail length of this species (referred to as the “costly signal hypothesis”). Although resident species on average have shorter tails than migrant species (Hasegawa et al. 2016), another resident congener with a shallowly forked tail, *Hirundo nigrita*, also uses nests above water, unlike the remaining six resident congeners (i.e. “nest sites are closely associated with water”; Turner and Rose 1994; also see Fig. 1), suggesting the importance of roosting location. Urban house-dwelling Pacific swallows provide a rare opportunity to test whether cold wintering conditions, and the anthropogenic alternative, affect ornament expression. In other words, we focused here on whether the focal ornament is maintained because of the benefits and costs of their expression or is a selectively neutral, obsolete signal.

**Figure 2.**
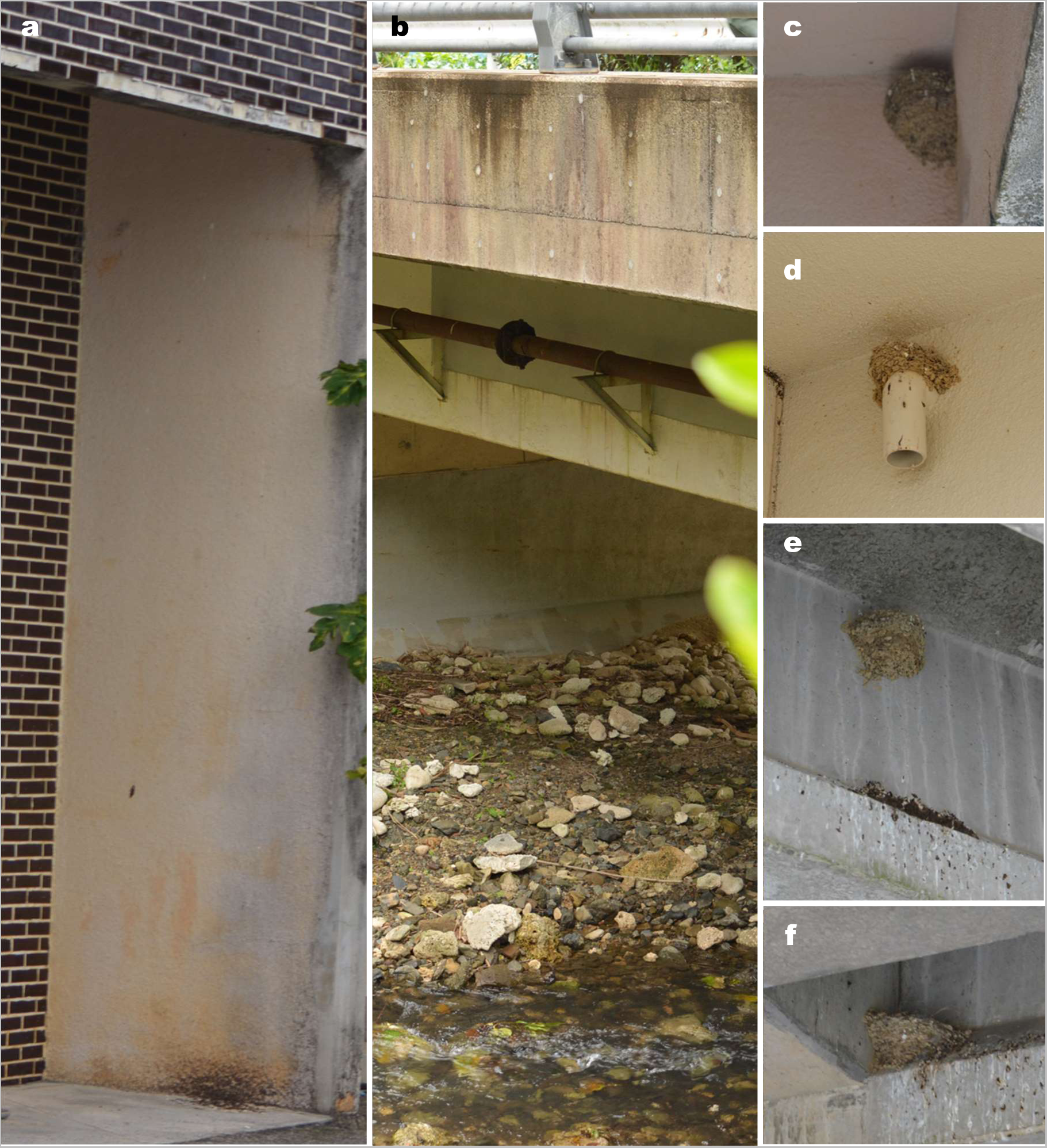
Examples of roosting sites of Pacific swallow on Amami Oshima Island: a) roosting site above ground observed from outside; b) roosting site under a bridge over a river observed from outside; c, d) roosting nests above ground; and e, f) roosting nests under bridges.

To examine ornament-specific pattern, besides outermost tail feather length we studied another sexually dimorphic plumage ornament, the red throat patch (Hasegawa et al. 2019). Given that a long outermost tail feather but not a colourful throat imposes a foraging cost (see the previous paragraph), the costly signal hypothesis (see above) predicts that individuals roosting under the eaves of houses will have longer outermost tail feathers compared to individuals roosting under bridges, whereas no detectable (or subtle) difference will occur in throat colouration. It should be noted that throat colouration is produced by eu- and pheo-melanin pigments, which also accompanies some maintenance cost (e.g., Arai et al. 2019; Hasegawa et al. 2019 for this species), but we here simply stress the importance of the foraging cost of the focal ornament, i.e., long tails, which directly reduce food intake required for somatic maintenance. To assess the importance of somatic maintenance, we also compared physiological state (i.e. body mass and plasma corticosterone level, here) between the two roosting locations with the prediction that individuals roosting under the eaves of houses would be in better physiological condition than those roosting under bridges (Diamond and Martin 2020). In particular, the plasma corticosterone level decreases with increasing local temperature (e.g. see Jenni-Eiermann et al. 2008 for barn swallows), and therefore we predicted that birds roosting under the eaves of houses would have lower plasma corticosterone levels than birds roosting under bridges. Finally, using an infrared thermometer, we examined whether the roosting sites under the eaves of buildings were in fact warmer than those under bridges over rivers. Because of the correlational nature of the current study, we discuss the observed pattern in the context of the costly signal hypothesis and alternative ecological and evolutionary explanations.

## MATERIAL & METHODS

### Capture and measurements

Capture survey was conducted in Amami Oshima Island in Kagoshima Prefecture, Japan (28°22′N, 129°29′E; total area 712.35 km^2^) from late-January to mid-February 2017–2020, mainly in the northern area, including a city (Amami-city, formerly known as Naze-city) and several towns (e.g. Tatsugo-cho, Yamato-son). Pacific swallows inhabit this island throughout the year, and few barn swallows breed or winter there (Hasegawa and Arai 2017a). Pacific swallows were captured in sweep nets while roosting at night. In these areas, swallows roost in and beside old nests under bridges over rivers or under the eaves of houses above the ground (Hasegawa and Arai 2017a). In the preceding year, 2016, we observed that some swallows used the eaves of houses above ground as nest sites, and therefore we carefully searched for roosts over land.

Upon capture, we marked each bird with metal rings with unique numbers (AC Hughes Ltd., Middlesex, UK), and collected blood samples (ca. 100 µL) from the brachial vein using heparinized capillary tubes within minutes of capture, although unfortunately we did not measure the exact time. We captured max. four individuals at once. The blood samples were immediately centrifuged to separate plasma (for the enzyme immunoassay, EIA, described below; Hasegawa et al. 2019) from red blood cells and other sediment (for sex identification; see Hasegawa and Arai 2017a). Blood samples were stored in an ice box until being returned to the hotel, and then at −20°C until being sent to the laboratory, where they were stored at −80°C (2017–2019) or −30°C (2020). MH measured the wing length, keel length, body mass, and outermost and central tail feather lengths. All lengths were measured to the nearest 0.01mm and body mass was measured to the nearest 0.1g (see Hasegawa et al. 2019 for detailed methods). At capture, we confirmed that all birds had completed the moult. We also collected a few throat feathers (ca. 10 feathers) to measure plumage colouration (see below) and traced the throat patch area by placing a transparent plastic sheet over the throat region. As described in Hasegawa et al. (2010), each bird’s throat patch was traced twice, and each plastic sheet was scanned and then the area of the red throat patch was measured. Throughout the study period, the repeatability of the measurements was moderate (repeatability = 0.65, *F_120,121_* = 4.50, *P* < 0.0001), perhaps due to the small variation in throat patch size compared to the barn swallow, in which we obtained high repeatability using the same methodology (repeatability > 0.80; Hasegawa et al. 2010). The mean of the two measurements was used for analyses.

We did not use data from 2016 in the current study because of the exceptional weather conditions in 2016, which affected the physiological state and survival of birds (Hasegawa and Arai 2017a). The sex of each individual was determined via molecular sexing (Hasegawa and Arai 2017a). Age differences in morphology, i.e. a potential confounding factor, are at best small in this species (Hasegawa et al. 2019; also see the Results section), although we conducted additional analyses using solely after-second-year (ASY) individuals, i.e. birds with previously attached rings after maturation, to exclude any possible age effect. This is because yearling individuals are often less-ornamented, at least in the congener, the barn swallow, *H. rustica* (e.g. Møller 1991; reviewed in Møller 1994; Turner 2006).

### Enzyme immunoassay

We separately analysed samples collected in each year. Blood plasma samples were analysed for corticosterone by EIA (Cayman #501320). Because preceding analysis demonstrated highly repeatable values between duplicates (repeatability = 0.98, *N* = 35, *F* = 80.05, *P* < 0.0001; Hasegawa et al. 2019), we performed a single analysis (i.e. did not duplicate) of the samples collected in 2017 to 2020. In 2017, the mean blood plasma volume per tube was 9.8 μL (range: 6–10 μL) and the standard curve fit the data well (*r* = 0.999). The sensitivity (80% maximum binding) and midpoint (50%) were 28.11 and 173.79 pg/mL, respectively, in 2017. Three samples fell outside the assay range of 8.2–5000 pg/mL in 2017 (all three >5000 pg/mL), two of which were re-assayed to estimate the corticosterone level. In the re-assay, the sensitivity (80% maximum binding) and midpoint (50%) were 72.22 and 322.77 pg/mL and the standard curve fit the data well (*r* = 0.93). Because some samples fell outside the assay range in 2017, we used 3 µL blood plasma for EIA in 2018 and the standard curve fit the data well (r = 0.997). The sensitivity (80% maximum binding) and midpoint (50%) were 21.07 and 115.25 pg/mL, respectively, in 2018. In 2019 and 2020, we used 5 µL blood plasma (except for one sample in 2019 from which we could not obtain 5 µL and used 4 µL instead) for EIA and the standard curve fit the data well (2019: *r* = 0.998; 2020: *r* = 0.982). The sensitivity (80% maximum binding) and midpoint (50%) were 21.12 and 109.83 pg/mL, respectively, in 2019. In 2020, the corresponding values were 43.42 and 161.32 pg/mL, respectively.

Unfortunately, we did not record time interval between capture and blood collection (see above), although this is necessary to statistically control for differential handling time, which may affect the results of plasma corticosterone levels (e.g., Jenni-Eiermann *et al*., 2008). Because we captured a maximum of four individuals before collecting blood when they roost together, plasma corticosterone levels might be increased when we handled multiple birds. Consistent with this possibility, individuals roosting >2 birds had higher plasma corticosterone levels than the others (MCMCglmm: *N* = 59, sex: coefficient = 0.08, 95%CI = –0.38, 0.52, *P_MCMC_* = 0.73; group roosting: coefficient = 0.62, 95%CI = 0.18, 1.10, *P_MCMC_* < 0.01). Nevertheless, the analysis excluding individuals roosting in group, all of which were roosting under bridges, revealed a significant relationship between roosting location and plasma corticosterone level (MCMCglmm: *N* = 40, sex: coefficient = 0.22, 95%CI = –0.24, 0.68, *P_MCMC_* = 0.37; roosting location: coefficient = –0.86, 95%CI = –1.41, –0.33, *P_MCMC_* < 0.001), indicating that the effect of roosting location shown in the Result section cannot be explained by group roosting alone. In fact, Pacific swallows roosting under bridges tended to roost in group when ambient temperature was low (MCMCglmm: *N_total_* = 63, *N_individual_* = 57, coefficient = –2.81, 95%CI = –4.09, –1.56, *P_MCMC_* < 0.001; Fig. S1), and hence the high plasma corticosterone level found in group roosting individuals (see above) might be partially due to cold weather conditions, although we could not demonstrate the relative importance of handling time and weather conditions from this analysis, which is beyond the scope of the current study.

Likewise, some roosting sites under bridges are difficult to access (e.g., due to differential water depth, 0-1m, and distance to our car, <100m) and thus might take longer times until blood collection than others. However, our results are inconsistent with this possibility, because there was no detectable effect of the roosting site on plasma corticosterone levels among birds captured under bridges with and without controlling for group roosting (MCMCglmm: both: *N* = 58, repeatability < 0.01; Fig. S2), indicating that not accessibility but common location (i.e., “under bridges” vs “under the eaves of houses”) explains the results (see Results). It should also be noted that roosting sites under the eaves of the houses also differ their accessibilities, because some roosting sites were located in the upper (2nd to 5th) floors and others were apart from parking site (but individuals still exhibited similar physiological state; see Results).

### Plumage reflectance measurement

In the laboratory, we piled five feathers on a piece of white paper so that the perimeters of the feathers coincided (Hasegawa et al. 2010). We obtained two feather samples from each bird. As in a previous study of the barn swallow *Hirundo rustica* (Hasegawa et al. 2022), we measured the reflectance of the feather samples using an Ocean Optics Jaz spectrometer with illumination provided by a PX-1 pulsed xenon lamp (Dunedin, FL, USA). We used two optical fibres with a collimating lens: the illuminating fibre was positioned at 90° to the horizontal surface of the feather samples to produce a small illumination spot (<1 mm in diameter), and the collecting fibre was set at 45°. Each sample was measured twice, replacing the sample between measurements, and averaged. From the generated spectra, we used the ‘TCS’ function in the R package “pavo” to calculate four colour measures for each feather sample within the tetrahedral colour space (Maia et al. 2013). Each colour can be described by a vector and represented by θ (horizontal angle), φ (vertical angle), and rA (achieved chroma; the proportion of that maximum achieved by the colour point), with θ and φ reflecting visible and ultraviolet hues, respectively. Increasing θ values indicate less-red colouration in the current study. A verbal description of φ cannot easily be provided, because it is beyond the trichromatic human visual sense. The repeatabilities of these measures throughout the study period were moderate (θ: repeatability = 0.57, *F_120,121_* = 3.71, *P* < 0.0001; φ: repeatability = 0.65, *F_120,121_* = 4.72, *P* < 0.0001; rA: repeatability = 0.63, *F_120,121_* = 4.43, *P* < 0.0001). The mean of the two measurements taken from two different feather samples per bird was used for analyses.

### Roosting site temperature

During 2023 wintering season (1–6 February), the temperature of the roosting sites was measured using an infrared thermometer (Testo 835-H1, Germany). Not to be confounded by the presence of birds, we measured the substrate temperature above empty nests or perching spots (i.e., not roosting spots with birds). We took only one measurement for each roosting site (i.e., only one recording per house or per bridge). To control for daily and hourly fluctuations of air temperature, we used air temperature in Naze city in the same hour (Japan Meteorological Agency 2023). In total, we collected data from 25 roosting sites (15 sites under bridges and 10 sites under eaves of houses), all of which were roosting sites where Pacific swallows were captured in the past, i.e., all data were collected from their actual roosting sites.

### Statistics

The phenogram, which represents a phylogenetic tree with time on the x-axis and a phenotypic trait on the y-axis, showing estimated evolutionary change in the trait over time, was computed using the function “phenogram” in the R package “phytools” (Revell 2012). Differences in morphologies and plumage colouration (see above) between two roosting locations were studied using a Bayesian linear mixed-effects model while controlling for sex. The identity of the birds and study year were used as random factors to account for repeated measurements of individuals across years and the potential year effect, respectively (note that we did not include the exact capture site to avoid complicating the model; this may not be problematic, as we conducted the survey across many sites on a relatively small island; see above). For this purpose, we used the ‘MCMCglmm’ function in the R package ‘MCMCglmm’ (Hadfield 2010). Because Pacific swallows moult in late summer (Turner and Rose 1994), individuals have a different set of phenotypic traits each year. Interactions between roosting location and sex on measurements were removed because all were nonsignificant (i.e. *P_MCMC_* > 0.05). We ran the analysis for 1,400,000 iterations, with a burn-in period of 600,000 and a thinning interval of 80 for each tree. The reproducibility of the MCMC simulation was confirmed by calculating the Brooks–Gelman–Rubin statistic (Rhat), which must be < 1.2 for all parameters (Kass et al. 1998). Mean coefficients and 95% credible intervals (CI) are shown. We repeated the analysis using only ASY individuals, although we could not test the interaction terms in these models due to the small sample size. Lastly, to test for age and year effects, we focused on individuals repeatedly captured in consecutive years at the same roosting locations (in this case, all used roosts under bridges), and conducted a paired t-test in relation to age (t, t+1) or year (favourable, unfavourable). When we captured the same individuals in more than two years, we used the first two years to avoid confounding with senescence (Møller and de Lope 1999). “Favourable” and “unfavourable” years were determined based on physiological conditions, body mass and plasma corticosterone level (unfavourable: 2018, 2020; favourable: 2017, 2019; see Fig. S2), although this classification could not distinguish between immediate and long-term effects of the roosting environment on physiological conditions (note that body mass had no detectable relationship with body size, measured as keel length, at least in this population; Hasegawa et al. 2019). Using these individuals, we also estimated between-year repeatability, i.e. the upper limit of heritability (Lessells and Boag 1987; Falconer and Mackay 1996). When analysing the binary response variable with fixed effects using MCMCglmm with “categorical” error distribution, we used a Gelman prior for fixed effects after standardizing each fixed effect with the residual variance fixed at 1 (see Hasegawa & Arai, 2021 for the rationale and details). Difference in the surface temperature of roosting sites was analysed using a linear model, and air temperature was included as a covariate (see the preceding section). All statistical analyses were conducted using R software (ver. 4.0.0; R Core Team 2020).

## RESULTS

### Roosting locations

Of a total of 126 individuals captured during the study period, 110 were roosting under bridges over rivers and the remaining 16 were roosting under the eaves of houses (binomial test, *P* < 0.0001). Of 16 individuals that were repeatedly captured across years, 14 consistently used the same roosting locations. Of the remaining two individuals, one changed its roosting location from under the eave of a house in the first year to under a bridge in the next year, while the other individual showed the opposite pattern. Below, we present analysis of all individuals with morphological measurements (*n* = 121). The detailed sample size and its composition for each analysis are shown in tables.

### Morphology

After statistically controlling for sex differences, we found no detectable difference in wing or keel length between the two roosting location types, i.e. under bridges and under the eaves of houses (Table 1; Fig. 3a,b). Likewise, we found no detectable difference in central tail feather length between the two roosting locations, although we observed reversed sexual dimorphism in this case, with males having shorter central tail feathers than females (Table 1; Fig. 3c). In contrast, after controlling for sex, individuals roosting under the eaves of houses had longer outermost tail feathers than birds roosting under bridges (Table 1; Fig. 3d). This difference is small relative to the mean outermost tail feather length (i.e. ca. 1% of mean tail length, 56.44 mm) but is more than 40% of the estimated sex difference (Table 1). Qualitatively similar results were obtained in the analysis of ASY (after-second-year) individuals alone (Table 1), although the sex difference in wing length was marginal in this comparison.

**Figure 3.**
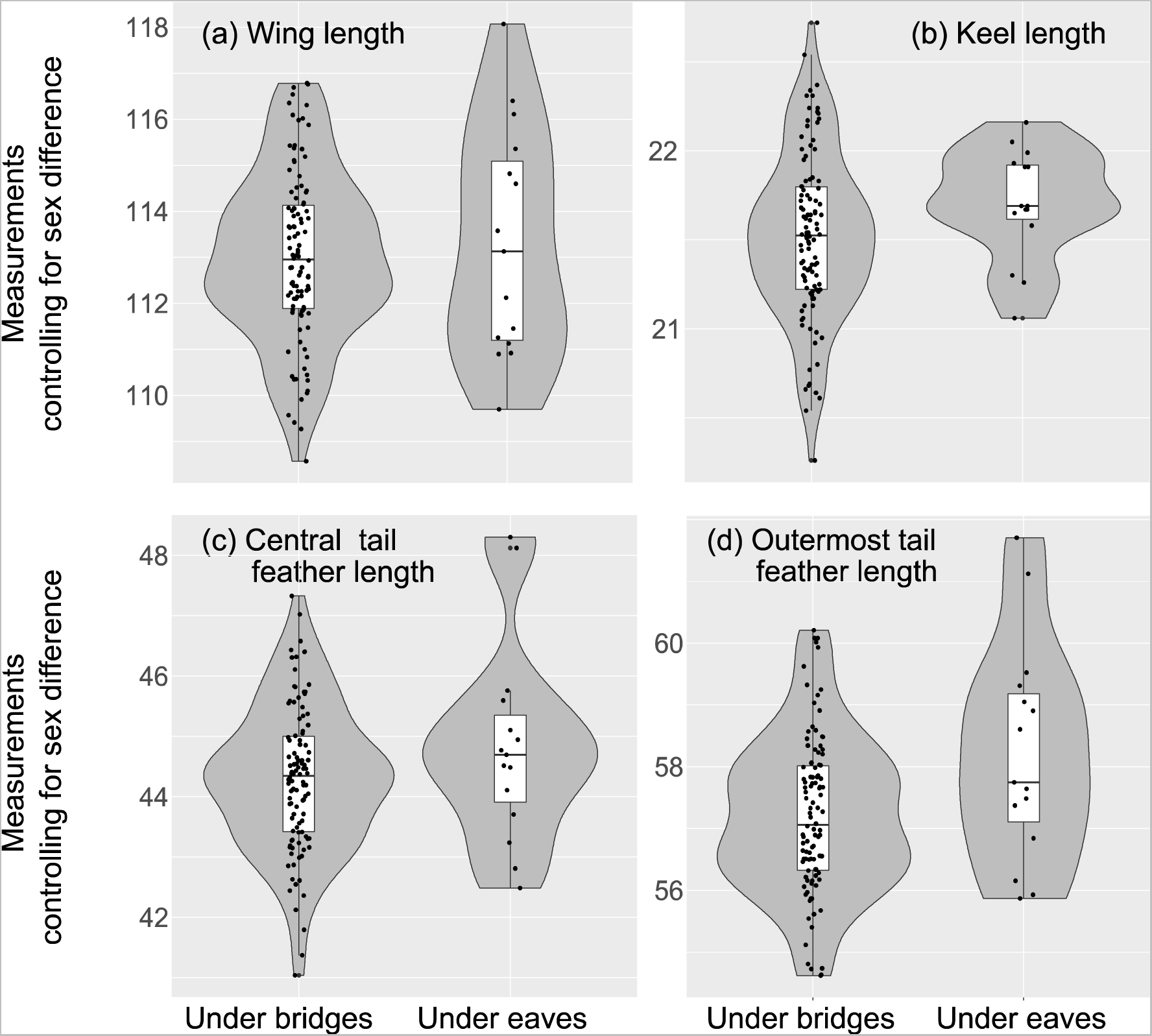
Violin plots of (a) wing length, (b) keel length, (c) central tail feather length, and (d) outermost tail feather length, all measured in millimetres, in relation to roosting location after controlling for sex differences (i.e. subtracting the estimated sex difference from female values) in the Pacific swallow during the 2017–2020 wintering seasons on Amami Oshima Island. Each measurement is standardised for each year with zero mean and unit variance to control for annual differences. Boxplots within violin plots indicate the median values and the first and third quartiles of the data (whiskers range from the lowest to the highest data points within 1.5 × the interquartile range of the lower and upper quartiles, respectively). The data points are shown as circles. See Table 1 for the formal statistics and sample sizes.

**Table 1.** Bayesian linear mixed-effects models for predicting each morphological measurement (i.e. wing length, keel length, central tail feather length, and outermost tail feather length) in Pacific swallows during the 2017–2020 winter periods.

| Variable | Wing length | Keel length | Central tail feather length | Outermost tail feather length |
| --- | --- | --- | --- | --- |
| All ages ($N_{all} = 121$, $N_{individual} = 103$) | | | | |
| Sex (male – female: S) | **1.99 [1.21, 2.75]\*\*\*** | **0.70 [0.52, 0.87]\*\*\*** | **–0.67 [–1.15, –0.16]\*** | **1.80 [1.27, 2.32]\*\*\*** |
| Roosting location (eave – bridge: R) | 0.34 [–0.64, 1.27] | 0.13 [–0.11, 0.35] | 0.34 [–0.29, 0.98] | **0.76 [0.07, 1.49]\*** |
| Interaction (S×R) | — | — | — | — |
| ASY alone ($N_{all} = 34$, $N_{individual} = 26$) | | | | |
| Sex (male – female: S) | 1.39 [–0.21, 3.03]† | **0.73 [0.41, 1.04]\*\*\*** | **–1.33 [–2.28, –0.39]\*\*** | **1.40 [0.28, 2.49]\*** |
| Roosting location (eave – bridge: R) | 0.80 [–1.24, 2.86] | 0.31 [–0.10, 0.70] | 1.19 [–0.10, 2.40] † | **1.43 [0.05, 2.78]\*** |
| Interaction (S×R) | NA | NA | NA | NA |

Mean coefficients and 95% credible intervals (CI) are shown.
Individual ID and study year were included as random effects.
All interaction terms for all ages were nonsignificant ( $P_{MCMC} > 0.05$ ) and thus were removed from the final models (note that we could not test the interaction term in ASY individuals alone due to the limited sample size, although each main factor, which includes more than five data points, could be successfully estimated).
Significant test results (i.e. $P_{MCMC} < 0.05$ ) are shown in bold (\* $P_{MCMC} < 0.05$ ; \*\* $P_{MCMC} < 0.01$ ; \*\*\* $P_{MCMC} < 0.001$ ; also note † $P_{MCMC} < 0.10$ ).
$N_{males}$ and $N_{females}$ for all ages were 60 (8) and 61 (7), respectively (numbers in parentheses are individuals roosting under the eaves of houses over the
ground). Likewise, $N_{males}$ and $N_{females}$ for ASY alone were 20 (4) and 14 (2), respectively.

### Throat feather colouration

Two measures of throat patch colouration, θ and φ, were explained by sex but not by roosting location (Table 2), with males having more colourful plumage with lower θ and higher φ than females (Fig. 4a,b). This was also the case for rA, although the effect of roosting location was marginal (Table 2; Fig. 4c). No detectable difference in patch area was found between the sexes and between roosting locations (Table 2; Fig. 4d). Qualitatively similar results were obtained when we analysed ASY individuals alone (Table 2), although both sex and roosting location were far from significant in the analysis of rA and the sex difference in φ was marginal.

**Figure 4.**
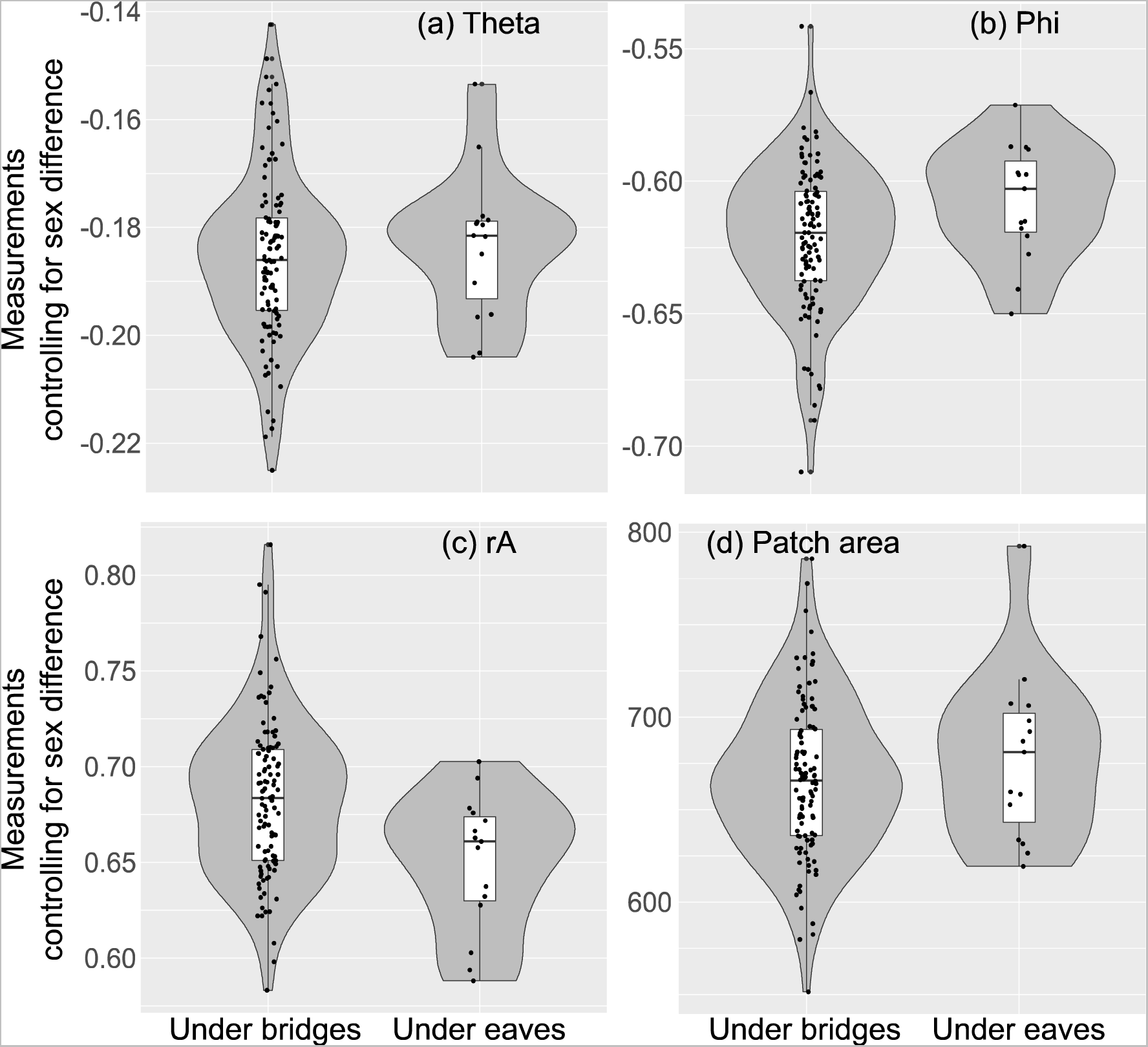
Violin plots of (a) theta, θ (angle, in radians), (b) phi, φ (angle, in radians), (c) rA (percent), and (d) patch area of red throat patch (mm^2^) in relation to roosting location after controlling for sex differences (i.e. subtracting the estimated sex difference from the female values) in the Pacific swallow during the 2017–2020 wintering seasons on Amami Oshima Island. Each measurement was standardised for each year with zero mean and unit variance to control for annual differences. Boxplots within violin plots indicate the median values and the first and third quartiles of the data (whiskers range from the lowest to the highest data points within 1.5× the interquartile range between the lower and upper quartiles). The data points are shown as circles. See Table 2 for the formal statistics and sample sizes.

**Table 2.** Linear mixed-effects models for predicting feather colour measurements (i.e. θ, φ, rA, and patch area) in Pacific swallows during the 2017–2020 winter periods.

| Variable | $\theta$ | $\phi$ | rA | Patch area |
| --- | --- | --- | --- | --- |
| All ages ($N_{all} = 121$, $N_{individual} = 103$) | | | | |
| Sex (male – female: S) | **–0.02 [–0.03, –0.02]\*\*\*** | **0.03 [0.02, 0.04]\*\*\*** | **–0.01 [–0.03, –0.004]\*** | 8.39 [–8.25, 25.24] |
| Roosting location (eave – bridge: R) | 0.00 [–0.01, 0.01] | 0.01 [–0.01, 0.02] | –0.02 [–0.04, 0.001]† | 10.25 [–12.28, 33.76] |
| Interaction (S×R) | — | — | — | — |
| ASY alone ($N_{all} = 34$, $N_{individual} = 26$) | | | | |
| Sex (male – female: S) | **–0.02 [–0.04, –0.01]\*\*\*** | 0.02 [0.00, 0.04]† | –0.01 [–0.03, 0.001] | –6.01 [–32.65, 22.26] |
| Roosting location (eave – bridge: R) | 0.00 [–0.01, 0.02] | 0.01 [–0.01, 0.03] | 0.00 [–0.03, 0.02] | 14.97 [–20.06, 49.54] |
| Interaction (S×R) | NA | NA | NA | NA |

Mean coefficients, and 95% credible intervals (CI) are shown.
Individual ID and study year were included as random effects.
All interaction terms were nonsignificant ( $P_{MCMC} > 0.05$ ) and thus were removed from the final models (note that we could not test the interaction term in ASY individuals alone due to the limited sample size, although each main factor, which includes more than five data points, could be successfully estimated).
Significant test results (i.e. $P_{MCMC} < 0.05$ ) are shown in bold (\* $P_{MCMC} < 0.05$ ; \*\* $P_{MCMC} < 0.01$ ; \*\*\* $P_{MCMC} < 0.001$ ; also note † $P_{MCMC} < 0.10$ ).
$N_{males}$ and $N_{females}$ were 60 (8) and 61 (7), respectively (numbers in parentheses are individuals roosting under the eaves of houses over the ground).
Likewise, $N_{males}$ and $N_{females}$ for ASY alone were 20 (4) and 14 (2), respectively.

### Physiological traits

Roosting site was a significant predictor of body mass (Table 3); individuals roosting under the eaves of houses were heavier than those roosting under bridges (Fig. 5a). Because the mean body mass during this period was 18.26, the estimated difference between the two locations (1.58) was roughly 9% of the mean body mass. Likewise, roosting location was a significant predictor of plasma corticosterone level (Table 3); individuals roosting under the eaves of houses had lower plasma corticosterone levels than those roosting under bridges (Fig. 5b). This difference (1.03) corresponds to 12% of the mean plasma corticosterone level (8.51, log-transformed value). There were no significant sex differences in these measures (Table 3). Qualitatively similar results were found when we analysed ASY individuals alone (Table 3).

**Figure 5.**
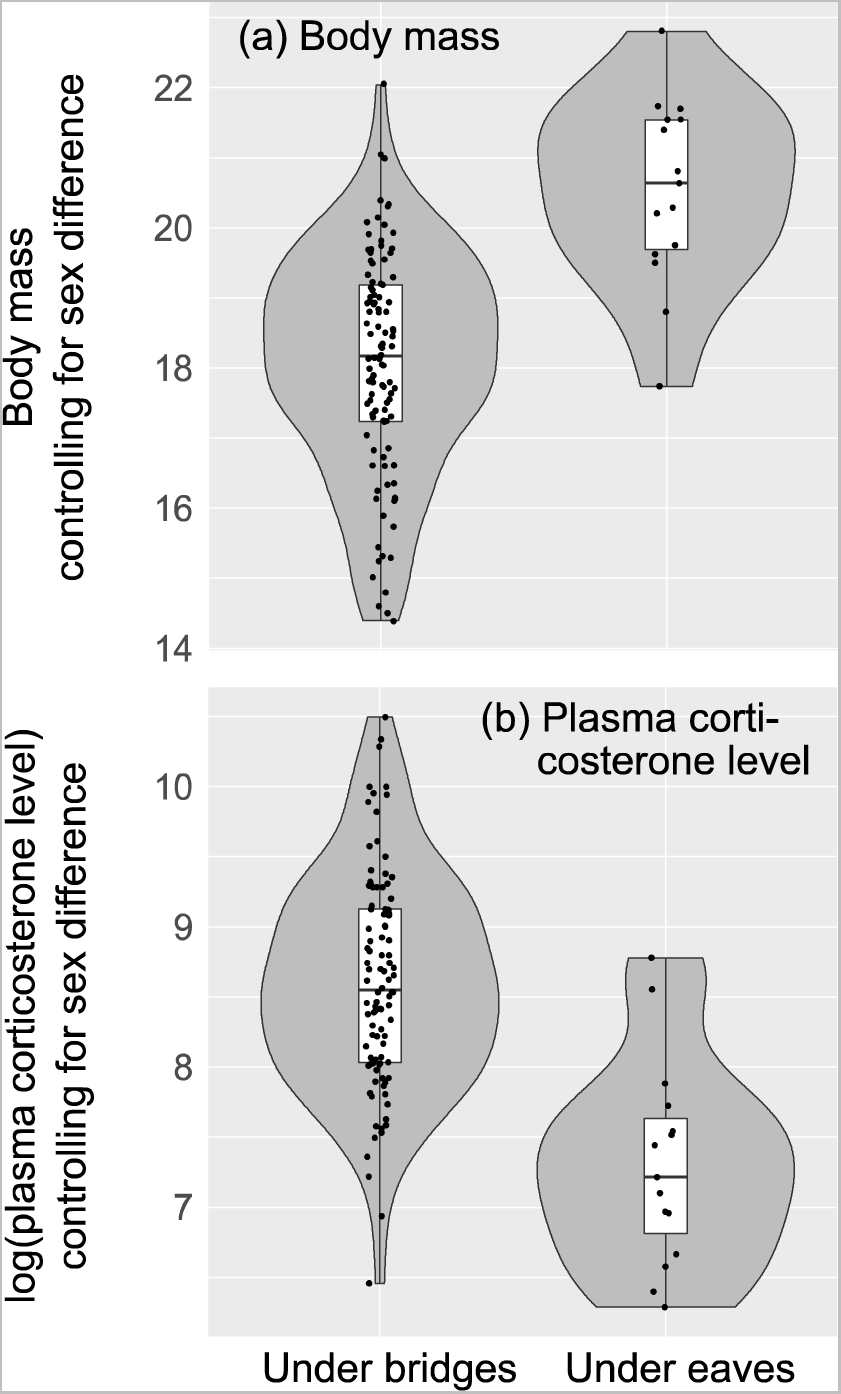
Violin plots of (a) body mass (g), and (b) plasma corticosterone level (pg/ml, log-transformed values) in relation to roosting location after controlling for sex difference (i.e. subtracting the estimated sex difference from female values) in the Pacific swallow during 2017–2020 wintering seasons on Amami Oshima Island. Each measurement was standardised for each year with zero mean and unit variance to control for yearly differences. Boxplots within violin plots indicate the median values and the first and third quartiles of the data (whiskers range from the lowest to the highest data points within 1.5× the interquartile range between the lower and upper quartiles). The data points are shown as circles. See Table 3 for the formal statistics and sample sizes.

**Table 3.**
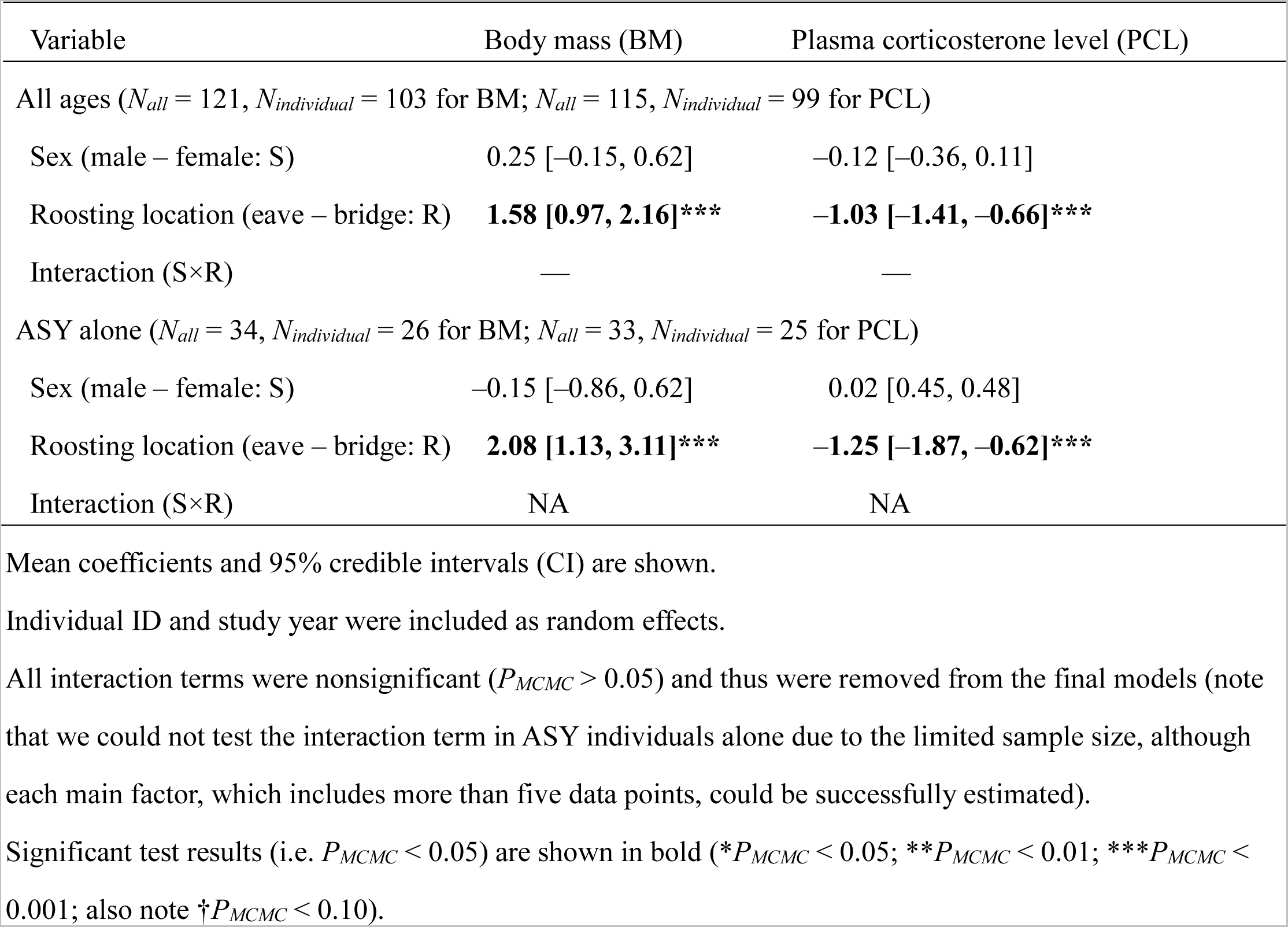
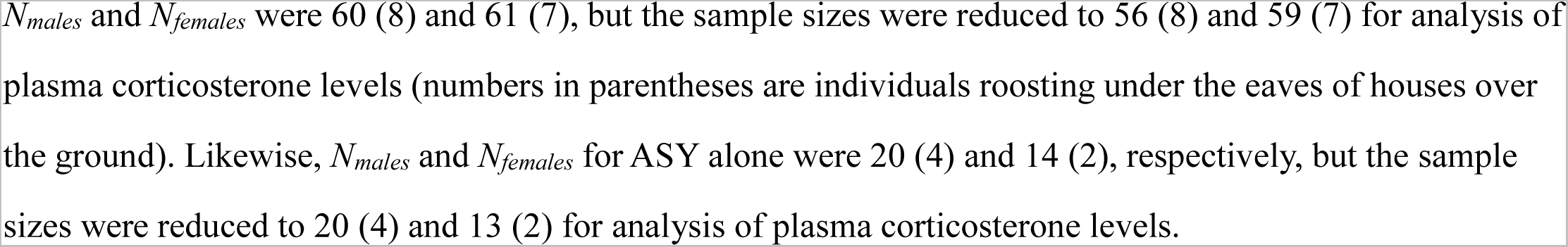
Linear mixed-effects models for predicting each physiological traits (body mass and plasma corticosterone level) in Pacific swallows during the 2017–2020 wintering seasons.

| Variable | Body mass (BM) | Plasma corticosterone level (PCL) |
| --- | --- | --- |
| All ages ($N_{all} = 121$, $N_{individual} = 103$ for BM; $N_{all} = 115$, $N_{individual} = 99$ for PCL) | | |
| Sex (male – female: S) | 0.25 [–0.15, 0.62] | –0.12 [–0.36, 0.11] |
| Roosting location (eave – bridge: R) | **1.58 [0.97, 2.16]\*\*\*** | **–1.03 [–1.41, –0.66]\*\*\*** |
| Interaction (S×R) | — | — |
| ASY alone ($N_{all} = 34$, $N_{individual} = 26$ for BM; $N_{all} = 33$, $N_{individual} = 25$ for PCL) | | |
| Sex (male – female: S) | –0.15 [–0.86, 0.62] | 0.02 [0.45, 0.48] |
| Roosting location (eave – bridge: R) | **2.08 [1.13, 3.11]\*\*\*** | **–1.25 [–1.87, –0.62]\*\*\*** |
| Interaction (S×R) | NA | NA |

Mean coefficients and 95% credible intervals (CI) are shown.
Individual ID and study year were included as random effects.
All interaction terms were nonsignificant ( $P_{MCMC} > 0.05$ ) and thus were removed from the final models (note that we could not test the interaction term in ASY individuals alone due to the limited sample size, although each main factor, which includes more than five data points, could be successfully estimated).
Significant test results (i.e. $P_{MCMC} < 0.05$ ) are shown in bold ( $*P_{MCMC} < 0.05$ ; $**P_{MCMC} < 0.01$ ; $***P_{MCMC} <$ $0.001$ ; also note $†P_{MCMC} < 0.10$ ).
$N_{males}$ and $N_{females}$ were 60 (8) and 61 (7), but the sample sizes were reduced to 56 (8) and 59 (7) for analysis of plasma corticosterone levels (numbers in parentheses are individuals roosting under the eaves of houses over the ground). Likewise, $N_{males}$ and $N_{females}$ for ASY alone were 20 (4) and 14 (2), respectively, but the sample sizes were reduced to 20 (4) and 13 (2) for analysis of plasma corticosterone levels.

### Age and year effects

In the comparison of individuals captured in consecutive years (year t and year t+1) in the same roosting locations, no detectable age effects were found in the ten measured traits (*N_individuals_* = 10, *N_total_* = 20, paired t-test, |*t*| < 1.89, df = 9, *P* > 0.09). Likewise, in the comparison of favourable and unfavourable years based on physiological conditions (see Methods; body mass: *t* = –6.05, df = 9, *P* < 0.0001; plasma corticosterone level: *t* = 6.61, df = 7, *P* < 0.0001), no significant yearly changes were found between the two years (i.e. unfavourable to favourable versus favourable to unfavourable) in any traits except throat colour rA (rA: paired t-test, *t* = 5.85, df = 9, *P* < 0.0001; all seven other traits: |*t*| < 1.71, df = 9, *P* > 0.12; Fig. 6). When we examined between-year repeatability, i.e. the upper limit of heritability, using these individuals, morphological variables showed moderate to high repeatabilities (wing length: rep = 0.84, *F_9,10_* = 11.37, *P* < 0.001; keel length: rep = 0.92, *F_9,10_* = 23.23, *P* < 0.0001; central tail feather length: rep = 0.80, *F_9,10_* = 8.95, *P* = 0.001; outermost tail feather length: rep = 0.67, *F_9,10_* = 4.99, *P* < 0.01), whereas throat colouration and physiological traits showed low to moderate repeatabilities (θ: rep = 0.81, *F_9,10_* = 9.31, *P* < 0.001; φ: rep = 0.28, *F_9,10_* = 1.78, *P* = 0.19; rA: rep = –0.61, *F_9,10_* = 0.24, *P* = 0.98; patch size: rep = 0.68, *F_9,10_* = 5.21, *P* < 0.01; body mass: rep = –0.28, *F_9,10_* = 0.56, *P* = 0.80; plasma corticosterone level: rep = –0.08, *F_7,8_* = 0.85, *P* = 0.58).

**Figure 6.**
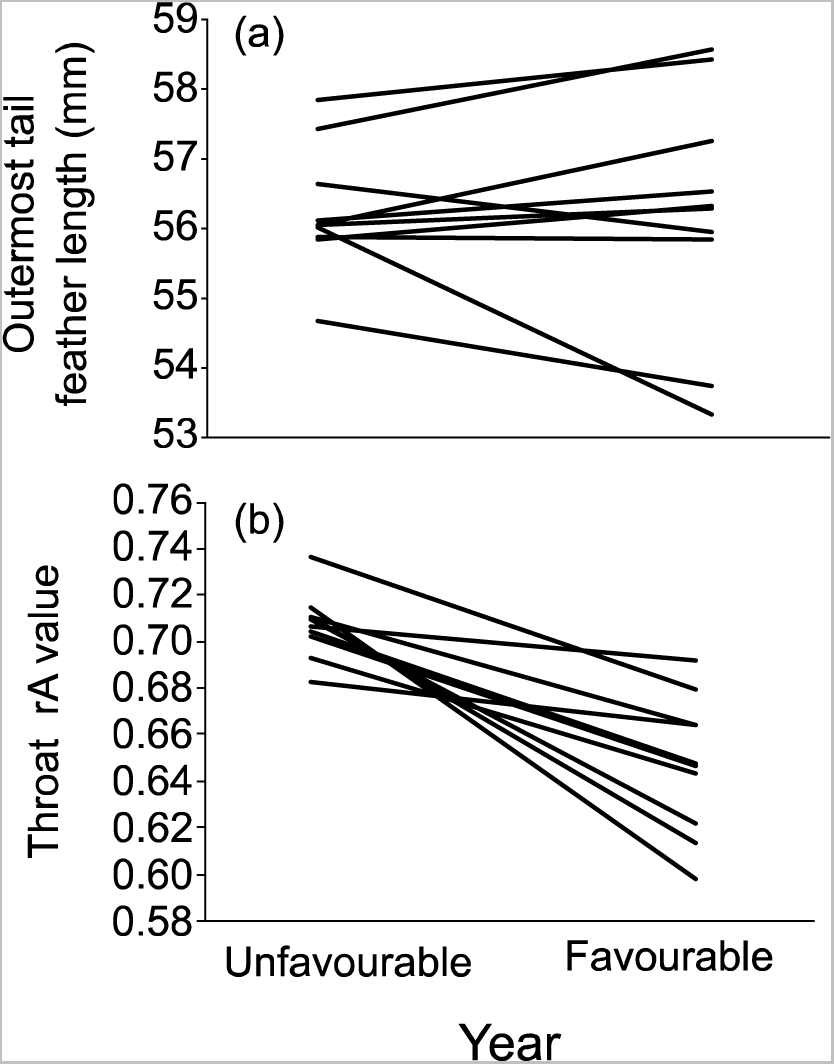
Pair-wise comparisons of (a) outermost tail feather length and (b) throat colour rA between unfavourable and favourable years for Pacific swallows captured in consecutive years at the same roosting locations. Lines connect observations for a given individual (both panels: *N_individuals_* = 10; see text for detailed information).

### Roosting site temperature

During 2023 wintering season, the substrate surface temperatures of the 25 roosting sites were measured using an infrared thermometer (n = 10 for roosting sites under the eaves of houses and n = 15 for roosting sites under bridges). After controlling for the significant effect of ambient air temperature (see Methods), we found that the roosting sites under the eaves of houses was approximately 1.6 °C higher than that of the roosting sites under bridges (Table 4; Fig. 7).

**Figure 7.**
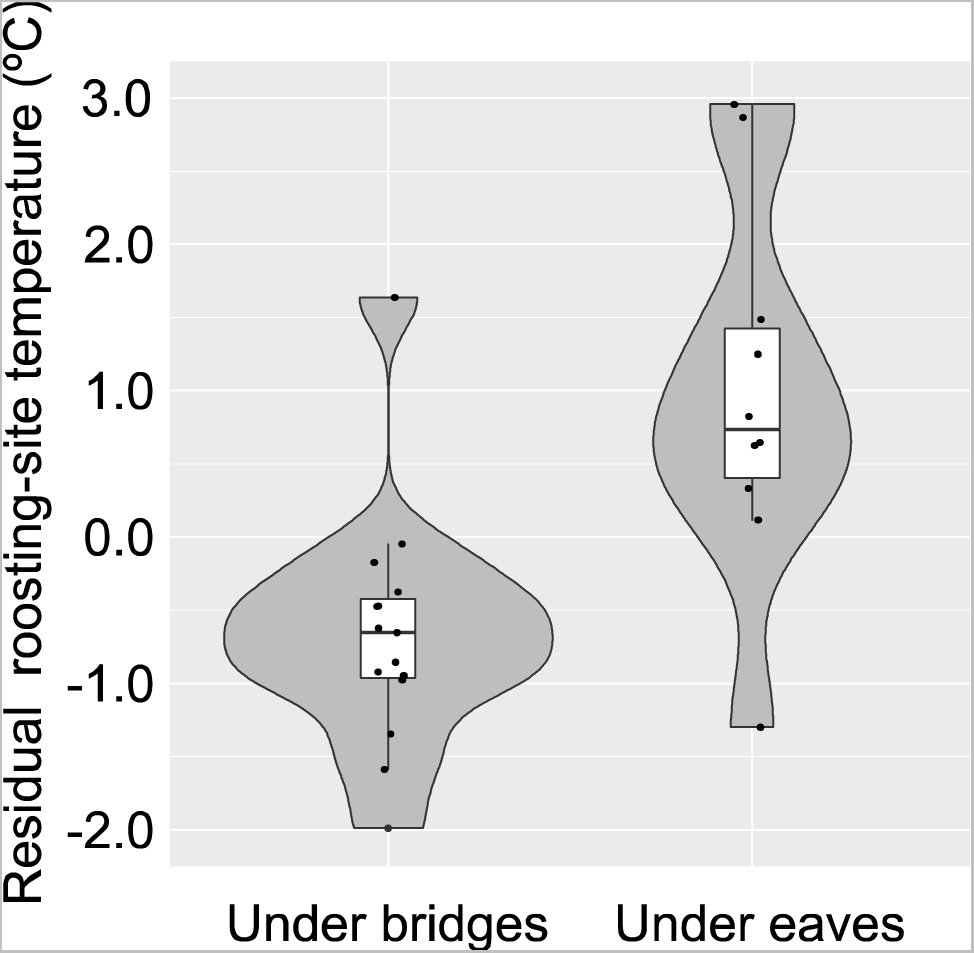
Violin plots of substrate surface temperature (°C) in relation to roosting location after controlling for daily fluctuation of air temperature during 2023 wintering seasons on Amami Oshima Island. Boxplots within violin plots indicate the median values and the first and third quartiles of the data (whiskers range from the lowest to the highest data points within 1.5× the interquartile range between the lower and upper quartiles). The data points are shown as circles. See Table 4 for the formal statistics and sample sizes.

**Table 4.** A linear model for predicting the surface temperature of the roosting sites of the Pacific Swallows during the 2023 wintering season (*N* = 25).

| Variable | Coefficients ± SE | $t$ | $P$ |
| --- | --- | --- | --- |
| Ambient air temperature | **0.31 ± 0.13** | **2.33** | **0.030** |
| Roosting location (eave – bridge: R) | **1.67 ± 0.43** | **3.91** | **<0.001** |

Mean coefficients, SE, test statistics ( $t$ ) and $P$ -values are shown. Interaction term was nonsignificant ( $P = 0.75$ ) and thus were removed from the final models. Significant test results (i.e. $P < 0.05$ ).

## DISCUSSION

The main finding of the current study is that roosting locations explained outermost tail feather length in the Pacific swallow; individuals roosting under the eaves of houses had slightly but significantly longer outermost tail feathers than individuals roosting under bridges. Because we controlled for the effects of sex and age, these factors would not confound our findings. In addition, because there was no detectable relationship between roosting location and feather lengths, measured by wing length and central tail feather length, the observed relationship would not be due to general growth pattern of feathers. The current study is one of few studies showing a link between ornamentation and anthropogenic wintering site (reviewed in Sepp et al. 2020), as predicted from the benign environment (i.e., warmer roosts; Fig. 7), which consistent with the good physiological conditions of birds roosting under the eaves of houses. We discuss possible explanations for these findings below.

A simple explanation for the observed patterns is differential costs and benefits of ornamentation between the two roosting locations, as predicted by the costly signal hypothesis. Individuals roosting under the eaves of houses (i.e. warmer roosts) had ca. 10% better physiological condition (i.e. heavier body mass, lower plasma corticosterone level) than individuals roosting under bridges. Therefore, these healthier individuals could afford greater investment in outermost tail feather length, a common secondary sexual trait in this clade (e.g., Hasegawa and Arai 2020a,b, 2022) that is constrained by its high maintenance cost associated with impaired foraging ability (e.g. see the Introduction; also see interspecific patterns in Hasegawa et al. 2016, Hasegawa and Arai 2018, 2020a,b, 2022). In addition to (relaxed) selection against long outermost tail feathers (Hasegawa et al. 2019), phenotypic plasticity may affect differential outermost tail feather length between the two roosting locations. In the barn swallow, a congener of the Pacific swallow, experimentally tail-elongated individuals in the preceding year have shortened tails after moult, perhaps due to impaired foraging ability, providing feedback on their ornamentation (Møller 1989), although natural outermost tail feather length in the barn swallow is affected by several other factors, including mite abundance and wintering conditions (e.g. Møller 1990, 1991; reviewed in Møller 1994; Turner 2006). Relatively long (but still short compared to congeners) outermost tail feathers have limited production costs, but outermost tail feather length shows a pattern consistent with condition-dependent expression, perhaps in part due to such plastic responses (e.g. see fig. 2 in Hasegawa et al. 2019 for this species; see Møller 1994; Turner 2006 for reviews in the barn swallow), reinforcing the link between roosting location and outermost tail feather length. Genetic and plastic effects are not mutually exclusive and in fact both were reported at least in the barn swallow (Møller 1989). Their relative importance in this study system remains to be clarified in the future.

The relative importance of genetic and plastic effects would also depend on ornament type. Analyses of yearly changes in ornamentation suggest that the importance of phenotypic plasticity might be lower for outermost tail feather length than throat colour rA, as only throat colour rA was enhanced in favourable years (Fig. 6; see also the moderate repeatability of tail length). This ornament-dependent plasticity is predictable based on the dynamic colour change that occurs after moulting, strengthening the relationship between physiological conditions and plumage colouration, because seasonal colour change via depigmentation, stains and wear depends on individual quality (e.g. Arai et al. 2015, 2019, Safran et al. 2010 and citation therein). Some Pacific swallows depart their nest/roosting sites and join communal roost during moulting period (Hasegawa 2020), and thus the post-moulting “dynamic” colour change rather than the moulting environment may explain the ornament-type dependent linkage to roosting location, which contrasts with a tail ornament, outermost tail feather length, which is static after moulting. Therefore, although the two ornaments measured here, outermost tail feather length and throat colouration (rA), appear to be similarly related to roosting location at first glance (Table 2), their relationships with roosting locations might be differently controlled.

In addition to the environment-driven differential ornamentation discussed above, ornament-dependent roost occupation or ornament-dependent dispersal might contribute to the observed pattern. For example, a possible explanation is that well-ornamented individuals are better able to hold roosts under the eaves of houses. This is unlikely, at least for outermost tail feather length, because plumage colouration rather than outermost tail feather length is used for intrasexual competition in the genus *Hirundo*, or at least the model species, *Hirundo rustica* (e.g. Hasegawa et al. 2014; Hasegawa et al. 2016; reviewed in Hasegawa 2018). However, outermost tail feather length might still affect roosting patterns if long-tailed individuals can offset unusual or non-preferred roosting locations with their own attractiveness to attract mates to their roosts. In this case, social mate attraction is beneficial for subsequent reproduction (e.g. early breeding onset) as well as for social huddling that improves thermoregulation (Hasegawa and Arai 2017b), thereby strengthening the link between physiological condition and roosting location. Consistent with this sexual (or social) selection explanation, male Pacific swallows have a shorter, rather than longer, central tail feather length, in addition to longer outermost tail feathers, compared to females (Table 1), which is predicted from the hypothesis of sexual selection favouring “forked” tails. That is, this pattern cannot be explained by sex differences in the size and general growth pattern of feathers (Møller 1994; Hasegawa and Arai 2020b, 2021). Nevertheless, the observed pattern is unlikely to have been caused solely by ornament-dependent roost occupation, as physiological differences between roosting locations are much larger than the difference in ornamentation between roosting locations and thus it is difficult to explain differential conditions as a consequence of ornament-dependent roost occupation. Therefore, rather than directly affecting the roost occupation pattern, sexual (social) selection would be the factor favoring long tails, which in turn affects ornament expression in combination with the cost of the ornamentation (see the preceding paragraph). Unfortunately, in the current correlational study with limited data, we cannot test this idea and other possible explanations (e.g. confounding factors other than age and sex), which should be explored in future studies.

An alternative explanation to those discussed above is that forked tails evolve and are maintained through viability selection for aerodynamic function (e.g. Norberg 1994; Evans 1998; Buchanan and Evans 2000). This explanation was previously applied in hirundines to explain the variety of fork depth and thus outermost tail feather length at least at interspecific levels, as shallowly and deeply forked tails are thought to have different fitness optima across the continuum of aerial foraging niches (e.g. speed versus manoeuvrability; Buchanan and Evans 2000; Park et al. 2001). However, experiments undertaken to test this aerodynamic explanation are theoretically invalid because, through manipulation of tail length alone, they ignore co-evolving compensatory traits, making estimation of the cost of forked tails impossible (e.g., Hasegawa and Arai 2020a,b; 2022; Hasegawa 2022; also see Norberg 1994, p. 16) and their findings are inconsistent with macroevolutionary patterns in swallows. Recent macroevolutionary studies indicate that the intensity of sexual selection, but not the changing direction of sexual selection or aerodynamic ability during foraging, can explain the evolution of forked tails and deeply forked tails are costly in terms of efficient aerial foraging (e.g., Hasegawa and Arai 2020a, 2022). Although these macroevolutionary studies focused on general interspecific patterns rather than specific functions in the focal species, we found patterns consistent with the idea that sexual selection favours long tails in the Pacific swallow. As noted above, Pacific swallows exhibit slight sexual dimorphism in outermost tail feather length and reversed sexual dimorphism in central tail feather length, which is inconsistent with the hypothesis that a viability advantage favours long tails (Hasegawa et al. 2019; Hasegawa and Arai 2020a). In addition, adults showed longer outermost tail feather lengths than juveniles (and only adults have elongated tail “streamer” parts; M. Hasegawa unpublished data; Fig. S3; also see Figs. 1 and 3), further supporting a sexual function of forked tails during reproduction. This perspective is consistent with the fact that the Pacific swallow forages on larger prey than the barn swallow, a congener with long outermost tail feathers (Turner and Rose 1994), in which long tails are costly for foraging (see above). Different aerodynamic phenotypic adaptations to local microclimates, for example, Pacific swallows might require faster flight under the eaves of houses than under bridges, favouring the combination of larger body mass, i.e., high wing loading, with longer outermost tail feathers, are also unlikely, as long outermost tail feathers reduce flight speed (e.g., Park et al. 2001). Notably, differences in aerodynamic adaptation alone do not explain the difference in plasma corticosterone level, a common measure of physiological stress (Table 3), and thus is insufficient to explain the overall observed pattern. Together, these arguments support a sexual, rather than viability, advantage of forked tails that offsets the viability disadvantage of forked tails (Hasegawa et al. 2019); however, we could not completely exclude the possible role of divergent viability selection favouring different tail shapes between the two locations due to differential benefits, rather than differential costs, of forked tails without conducting manipulative experiments.

In conclusion, we found differential ornamentation in relation to roosting locations in a poorly-ornamented species, the Pacific swallow. The observed patterns indicate that reduced tail ornamentation, which is explained by physiologically demanding, cool roosting locations, can be somewhat exaggerated in favourable anthropogenic, warm roosts. In other words, the short tails of Pacific swallows would not be a selectively neutral, obsolete signal, which is consistent with the idea that these ornaments are maintained based on the benefits and costs of their expression. Previous studies of animal ornamentation have mainly focused on breeding periods, but wintering periods, during which somatic maintenance is critical, can be an important period differentiating ornamentation based on the local environment.

## ACKNOWLEDGEMENTS

We greatly appreciate all house owners who allowed us to study Pacific swallows in their houses. We also thank Dr Taku Mizuta for managing our field study. We appreciate Dr Nobuyuki Kutsukake and his lab members at Sokendai, and Dr Shumpei Kitamura and his lab members at Ishikawa Preferctural University. We thank Drs Yohei Terai, Eiji Tanaka, Tohru Ikeya, and Ichiro Tayasu for their technical support on laboratory experiments. MH was supported by a Research Fellowship of the Japan Society for the Promotion of Science (JSPS, 15J10000) and KAKENHI (JSPS, 19K06850).

---

**Figure S1.**
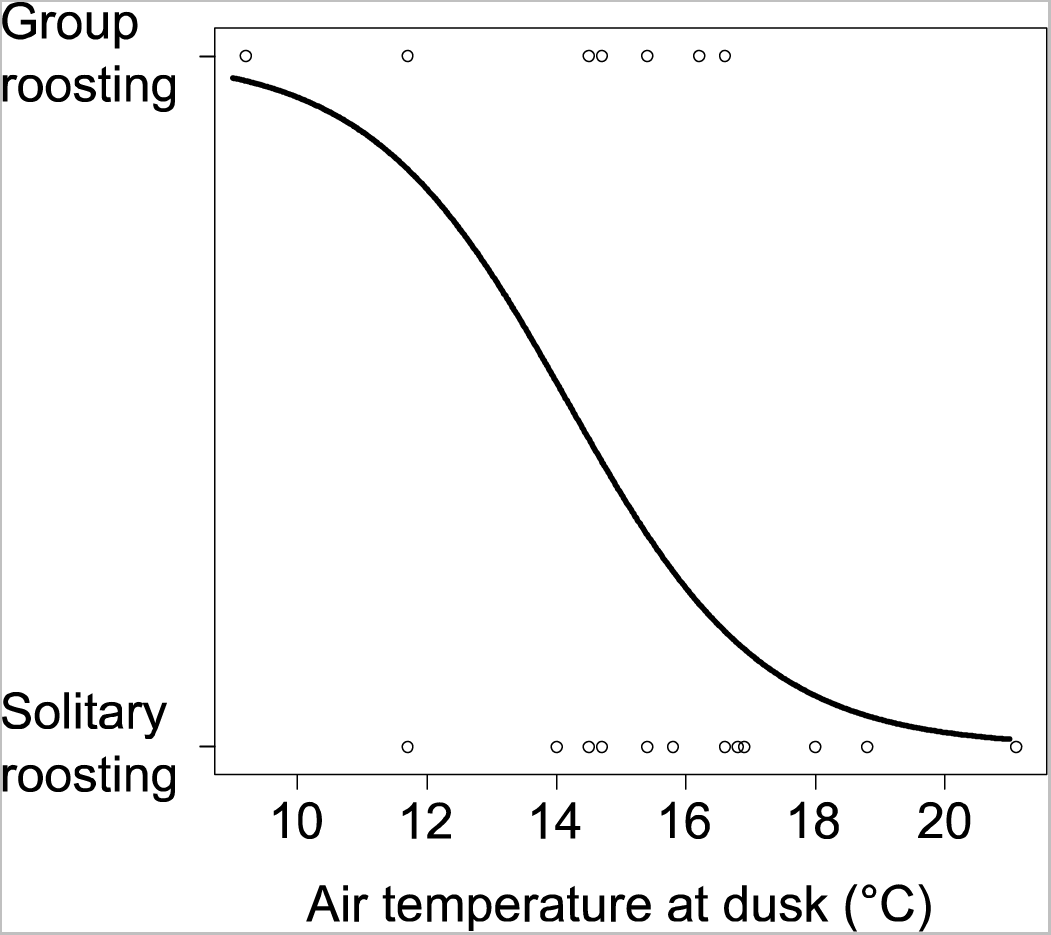
Relationship between air temperature at sunset and group/solitary roosting under bridges in the Pacific swallows wintering in Amami Oshima island. Air temperature at dusk is given by Japan Meteorological Agency (2023). Circles indicate the group or solitary roosting, and the line predicting the probability of group roosting is plotted (see text for statistics).

**Figure S2.**
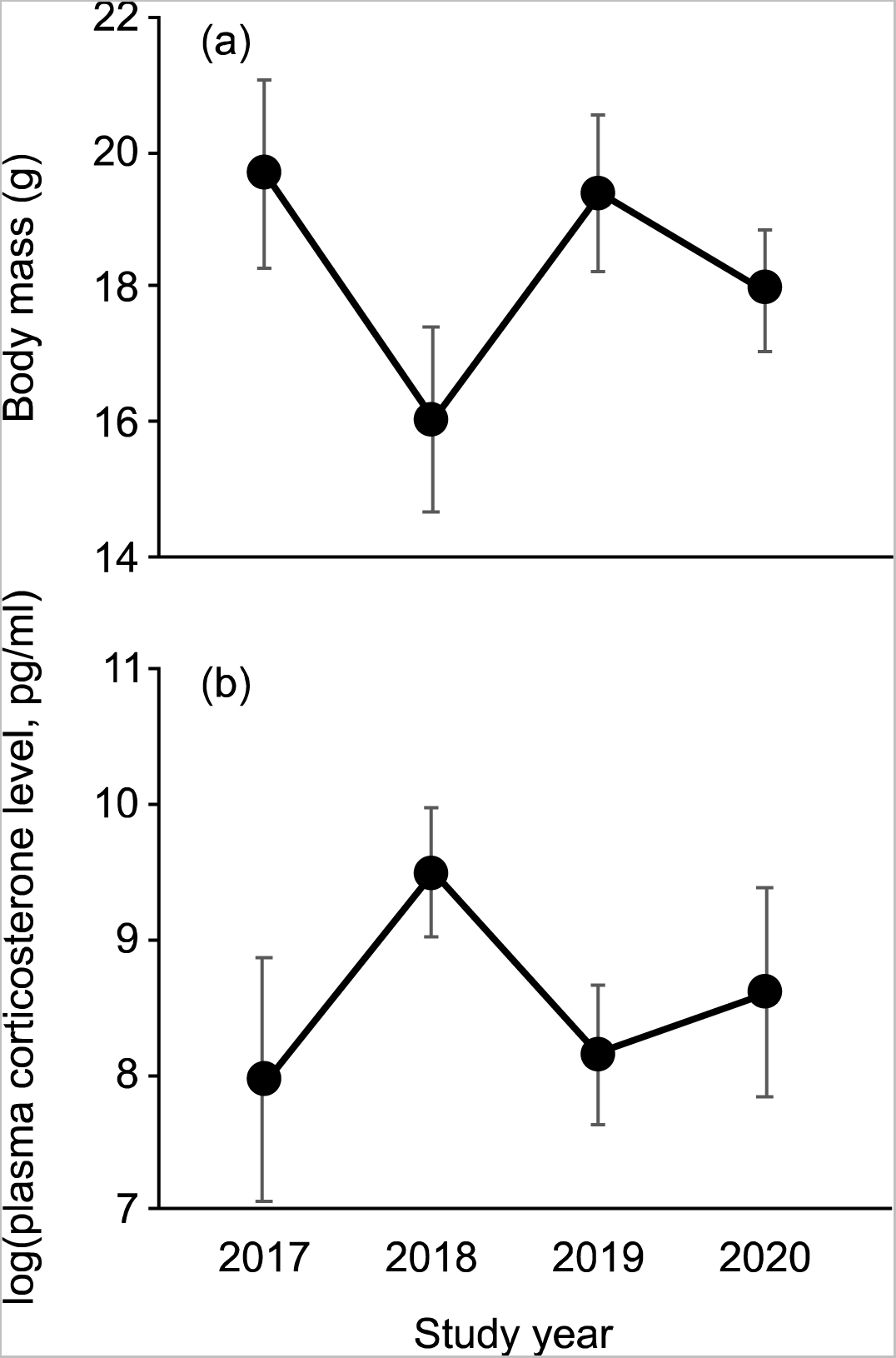
Yearly changes in (a) mean (± SD) of body mass and (b) plasma corticosterone level in the Pacific swallow during the 2017–2020 wintering seasons on Amami Oshima Island (see text for detailed information). From the four years of this study, the weather conditions of 2017 and 2019 were considered favourable, and 2018 and 2020 were considered unfavourable (see text).

**Figure S3.**
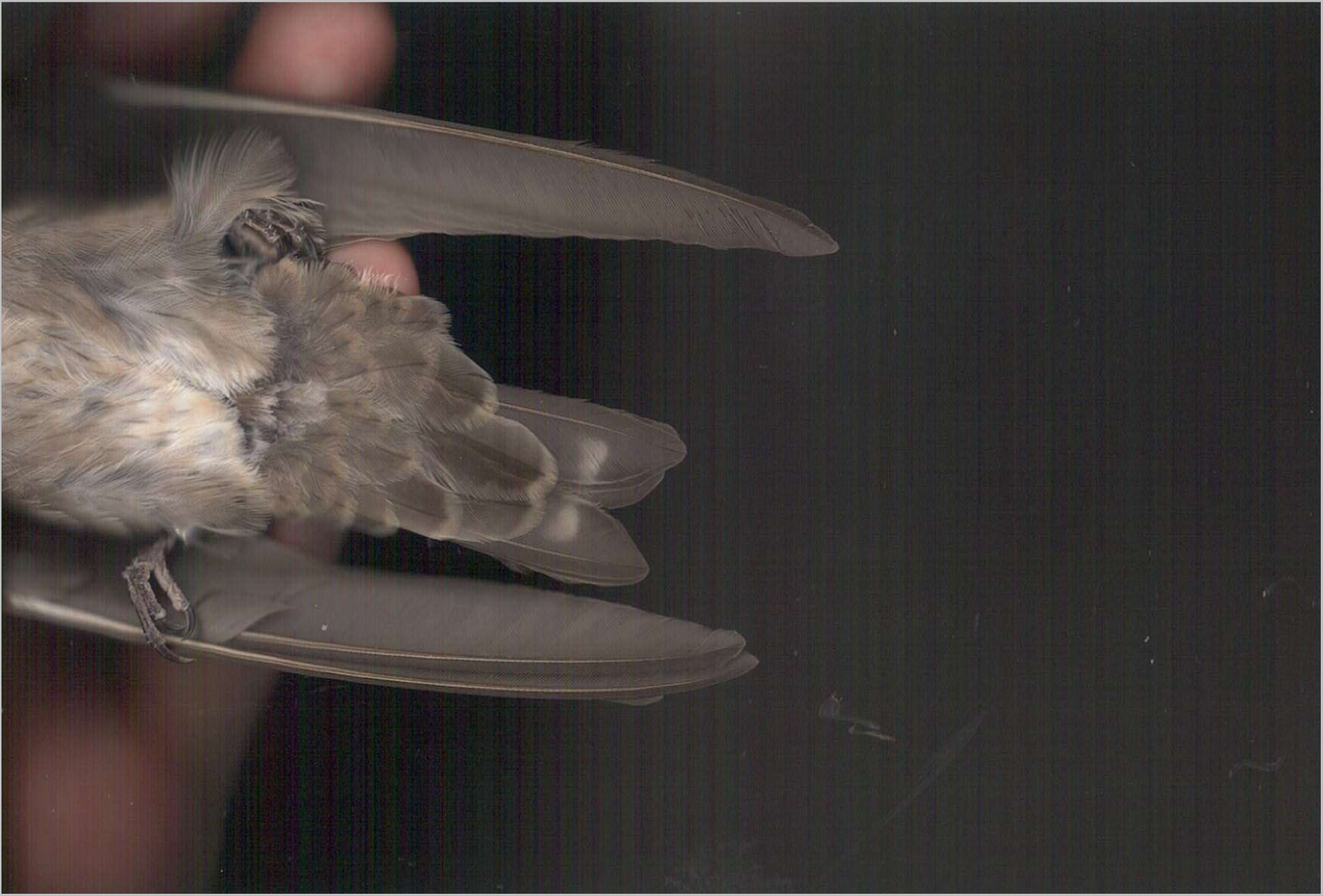
Juvenile Pacific swallow lacking the “streamer” part of the outermost tail feathers (cf. Fig. 1). In juveniles, mean central and outermost tail feather lengths ± SD (sample size) are 44.18 ± 0.80 (9) and 49.76 ± 1.46 (10), respectively.

